# A streamlined, cost-effective, and specific method to deplete transcripts for RNA-seq

**DOI:** 10.1101/2020.05.21.109033

**Authors:** Amber Baldwin, Adam R Morris, Neelanjan Mukherjee

## Abstract

RNA-sequencing is a powerful and increasingly prevalent method to answer biological questions. Depletion of ribosomal RNA (rRNA), which accounts for 80% of total RNA, is an extremely important step to increase the power of RNA-seq. Selection for polyadenylated RNA is a commonly used approach that excludes rRNA, as well as, important non-polyadenylated RNAs, such as histones, circular RNAs, and many long noncoding RNAs. Commercial methods to deplete rRNA are cost-prohibitive and the gold standard method is no longer available as a standalone kit. Alternative non-commercial methods suffer from inconsistent depletion. Through careful characterization of all reaction parameters, we developed an optimized RNaseH-based depletion of human rRNA. Our method exhibited comparable or better rRNA depletion compared to commercial kits at a fraction of the cost and across a wide-range of input RNA amounts.

## Introduction

RNA-sequencing (RNA-Seq) has become a widely accessible and powerful tool for answering questions in the field of biology resulting in a wealth of new insights. One major hurdle of RNA-seq library preparation is ribosomal RNA (rRNA) as it is highly abundant and makes up 80-90% of total RNA. To be cost-effective and achieve detection and quantification of less abundant transcripts, it is necessary to deplete samples of rRNA prior to library preparation. Many researchers enrich for polyadenylated transcripts using oligo-dT prior to library preparation to effectively deplete rRNA along with other nonpolyadenylated RNAs. Although oligo-dT selection is costeffective and accessible to most molecular biologists, there remain numerous drawbacks. It is not feasible for many samples including low quality or degraded FFPE and bacterial RNA. Oligo-dT selection will not capture the many important coding and non-coding non-polyadenylated transcripts (Cui et al., 2010; Guo et al., 2015; Sultan et al., 2014). Furthermore, it is far less effective at capturing precursor RNA important for investigating RNA metabolism (Mukherjee et al., 2017; Rabani et al., 2011; Zhao et al., 2014).

Current methods available for rRNA depletion predominantly fall into two categories, subtractive hybridization and RNaseH-mediated depletion (Figure 1). In subtractive hybridization, reverse complementary DNA probes with a 5’biotin moiety are hybridized to target rRNA. The RNA:DNA hybrid is then immobilized to streptavidin beads on a magnet and the transcripts remaining in the supernatant are used for downstream library preparation. This method has been developed and applied successfully for many samples and species (Culviner et al., 2020; Kim et al., 2019; Kraus et al., 2019; Su and Sordillo, 1998), but can become quite expensive with the cost of magnetic streptavidin beads required and synthesis of biotinylated oligos. This is especially true if multiple rounds of depletion are required to achieve higher efficiency (Kraus et al., 2019). RNaseH-mediated rRNA depletion is performed similarly in that reverse complementary DNA oligos (not biotinylated) are hybridized to rRNA to form RNA:DNA hybrids. The sample is then treated with RNaseH endonuclease which will specifically cleave RNA in the RNA:DNA, followed by a DNase step to remove remaining DNA oligos. This method is also highly efficient, and works well for low quality samples such as archived FFPE sections (Adiconis et al., 2013; Haile et al., 2019; Morlan et al., 2012).

**Fig. 1.**
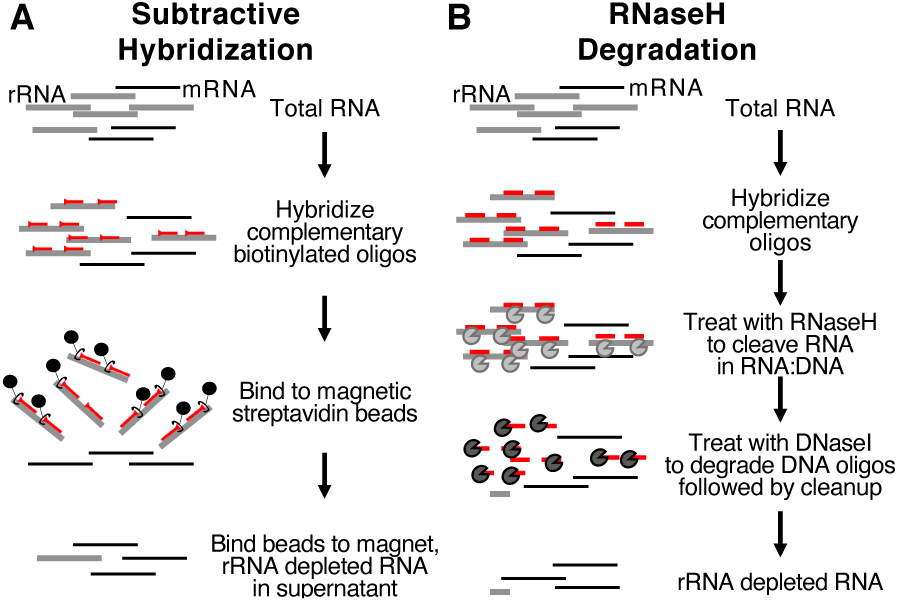
Current methodology for rRNA Depletion. A) rRNA from a total RNA sample is hybridized to biotin-linked reverse complementary oligos. The hybridized oligos are then bound to magnetic streptavidin beads to pull down the rRNA. After a heating step to release any nonspecifically bound transcripts, the supernatant containing the rRNA-depleted RNA fraction is collected and further purified by ethanol precipitation, bead cleanup, or column cleanup. B) rRNA from a total RNA sample is hybridized to reverse complementary DNA oligos to form RNA:DNA hybrid. The sample is then treated with RNaseH to cleave the RNA in the RNA:DNA hybrid, followed by DNaseI treatment to further cleave DNA oligos. The remaining rRNAdepleted RNA fraction is further purified by ethanol precipitation, bead cleanup, or column cleanup.

As the cost of sequencing continues to become less expensive, library preparation can make up a large portion of the price for an experiment. Commercially available kits offer ease of use and accessibility, but are more expensive than established “home-brew” methods that are easily performed in the average molecular biology lab (Table 1). Furthermore, commercial kits have their limitations, including RNA input amount which can be problematic for precious samples with very little RNA, as well as for applications requiring more than 5 ug of input RNA. As most details of these kits are considered proprietary, the kits are less flexible due to the lack of probe-design transparency. One commercially available kit, Illumina’s RiboZero, was considered the gold standard for rRNA depletion but is no longer available as a standalone kit. Additionally, labs working with less common organisms may not be able to find a commercially produced kit for rRNA depletion. Finally, researchers may want to deplete other highly abundant RNAs to improve depth and breadth of transcript detection and quantification which is not possible with many commercially available kits.

**Table 1.**
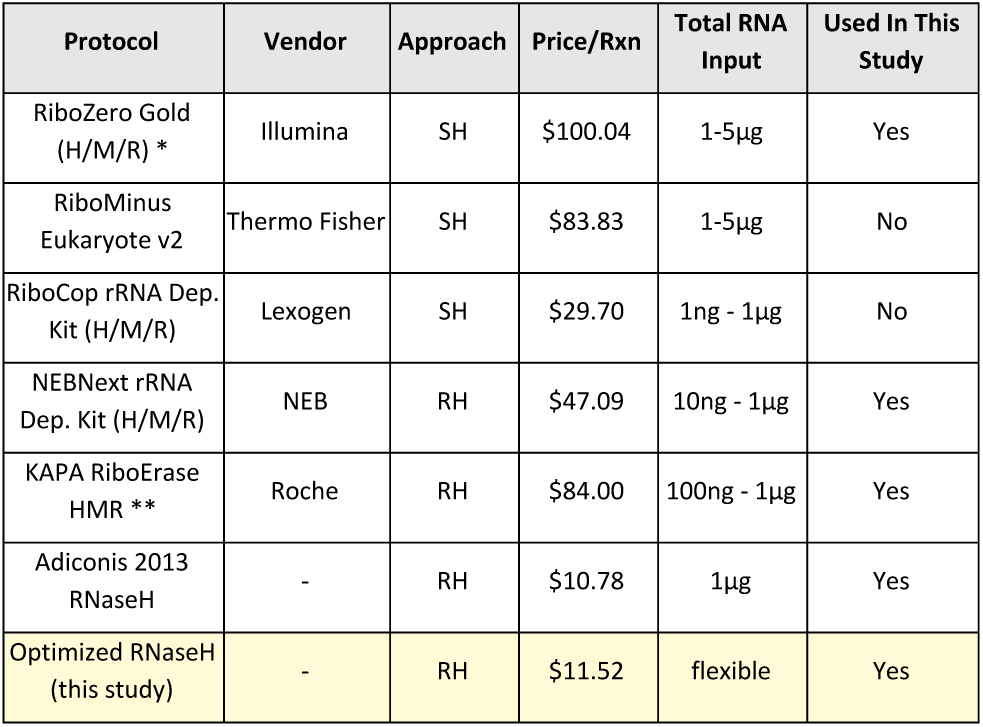
List price per reaction and total RNA input requirements for several commercially available human/mouse/rat rRNR depletion kits and one published method. SH = subtractive hybridization; RH = RNaseH-mediated. *lllumina Ri-boZero and RiboZero Gold (H/M/R) no longer available as a stand-alone probuct; Price reflects previous stand-alone product price per reaction. **KAPA RiboErase not available as a stand-alone product; Price reflects rRNA depletion plus library preparation kit total price per reaction.

Our goal was to optimize and describe a streamlined, costeffective, and flexible method for depleting rRNA and other abundant transcripts from human samples while maintaining minimal off-target depletion. Utilizing an RNaseH-mediated method, we have carefully characterized reaction conditions including reaction time, temperature, type of RNaseH enzyme, and DNA oligo:RNA input ratios. We compared the performance of our optimized method to that of popular commercially available kits. We also demonstrate that our method is applicable to a wide-range of input RNA amounts.

## Results

### Efficient but non-specific RNaseH-mediated rRNA depletion

We first sought out to apply a streamlined and costeffective ribosomal RNA depletion method similar to previously published methods using RNaseH. First, we hybridized 50-mer DNA oligos reverse complementary to human cytoplasmic and mitochondrial rRNA against total RNA isolated from human cell lines (H295R and HEK293). The samples were treated with E.coli RNaseH to cleave RNA in the RNA:DNA hybrid and further treated with E.coli DNaseI to clean up DNA oligos. To assess rRNA depletion, the resulting reactions and controls were run on an agarose gel. Substantial depletion of rRNA was only observed in the presence of both oligos and RNaseH (Figure 2A). Very modest fragmentation occurred in control samples, presumably due to the temperature and divalent cations required in the reactions. The DNA oligos appeared incompletely digested by DNaseI. Next, we assessed the specificity of depletion by performing RT-qPCR with primers to interrogate 18S rRNA (on-target) and ACT and GAPDH mRNAs (off-target). Compared to input RNA, we detected more 18S rRNA in samples treated with DNA oligos and DNaseI, but not RNAseH, indicating that incompletely digested oligos were serving as template for amplification and that the 18S rRNA depletion detected by RT-qPCR was an underestimation (Supplemental Figure 1). For this reason, all subsequent depletion samples were compared to the no-oligo control sample, which represents a reaction fragmentation control. Consistent with the gel electrophoresis, 18S rRNA was substantially depleted (Figure 2B). However, both ACTB and GAPDH mRNA were also depleted compared to no-oligo control, though not to the same extent as 18S rRNA. These experimental conditions resulted in a low accuracy (high on-target and medium offtarget) rRNA depletion.

**Fig. 2.**
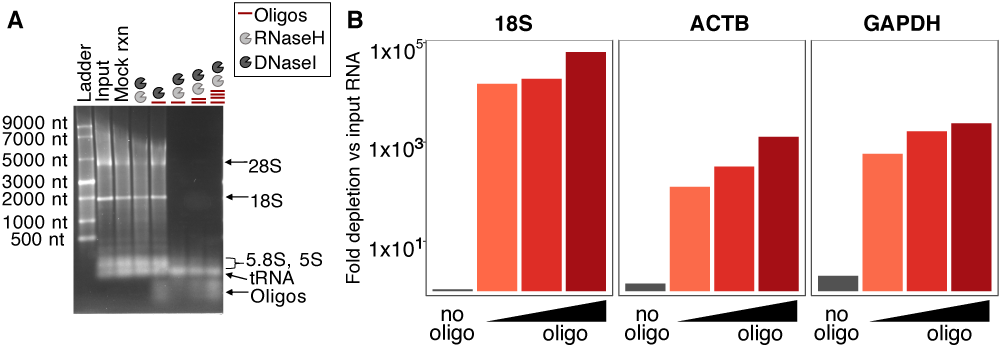
RNaseH-mediated rRNA and non-specific mRNA depletion. A) Nondenaturing 1.2% agarose gel depicting the following lanes from left to right: 1) ssRNA ladder; 2) total RNA input; 3) input with mock incubations; 4) RNaseH treatment and DNaseI treatment without oligos; 5) DNaseI and oligos only; and 6-8) increasing oligo:RNA mass ratio (1:1, 2:1, and 4:1; total RNA fixed at 1 µg) with RNaseH and DNaseI treatment. B) Fold depletion of 18S rRNA, ACTB, and GAPDH transcripts normalized to input total RNA by RT-qPCR performed on the RNA samples in panel A. Total RNA samples incubated with RNaseH and DNaseI without oligos (grey bar) or increasing oligo:RNA ratio (red bars) with RNaseH and DNaseI treatment. NEB E.coli RNaseH treatment for 1 hour at 37°C and NEB E.coli DNaseI treatment for 30 minutes at 37°C used for samples in this experiment.

### Optimization of rRNA depletion specificity

In order to maximize on-target and minimize off-target depletion, we focused on optimizing the RNaseH incubation time, type of RNaseH enzyme used in the treatment, and temperature of the reaction. While reducing incubation time while using E.coli RNaseH decreased off-target mRNA depletion, ontarget depletion also decreased (Figure 3A). We next introduced Hybridase Thermostable RNaseH, which has enzymatic activity at temperatures ranging from 45°C 95°C (Lucigen, 2019). We reasoned that a higher reaction temperature would result in higher hybridization specificity, leading to a higher accuracy of depletion. The Thermostable RNaseH reaction at 45°C yielded more on-target depletion of 18S rRNA and minimal to no off-target transcript depletion in comparison to E.coli RNaseH (Figure 3B). Increasing the Thermostable RNaseH reaction temperature to 65°C led to a dramatic increase in accuracy of on-target depletion while maintaining minimal off-target effects (Figure 3C). Varying the DNA oligo:input RNA ratio did not seem to have a large effect on accuracy of depletion by Thermostable RNaseH at the same temperature (Figure 3D). Taken together, our data indicate utilizing Thermostable RNaseH at 65°C for 10 minutes provided the best balance between depletion efficiency and specificity.

**Fig. 3.**
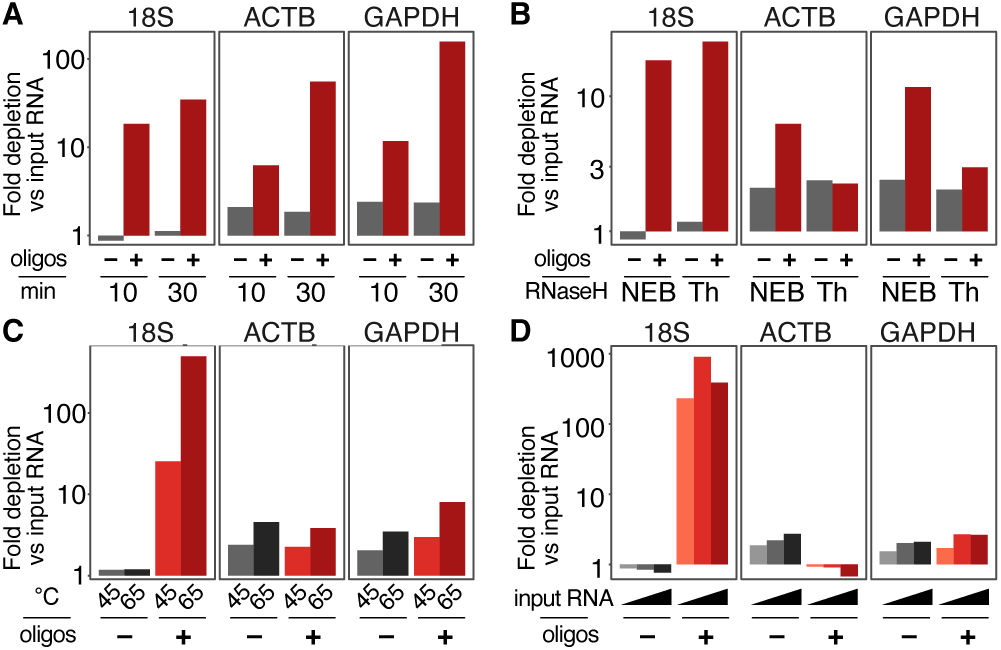
Optimizing the time, enzyme, temperature, and oligo concentration on rRNA depletion efficiency and specificity. For all experiments, fold depletion of 18S, ACTB, and GAPDH transcripts were normalized to input total RNA. Total RNA sample incubated with or without 1 µg of DNA oligos (grey and red, respectively). A) Varying reaction time using NEB E.coli RNaseH for 10 or 30 minutes at 37°C. B) Comparison of NEB E.coli RNaseH (NEB) and Lucigen Hybridase Thermostable RNaseH (Thermo) both with 10 minute incubations at 37°C and 45°C, respectively. C) Varying reaction temperature (45°C or 65°C) using Lucigen Hybridase Thermostable RNaseH for 10 minutes. D) Varying oligo:RNA ratio by increasing input RNA (1:1, 1:2, and 1:4; DNA oligos fixed at 1 µg) using Lucigen Hybridase Thermostable RNaseH for 10 minutes at 65°C.

### Comparison with multiple commercial rRNA depletion kits

Since our ultimate goal was to enable cost-effective rRNA depletion for RNA-seq, and given some of the caveats of RT-qPCR analysis, we next performed RNA-seq using the optimized parameters. We constructed duplicate RNAseq libraries using either our rRNA depletion approach, a previously published method (Adiconis), and commercially available kits. Although RiboZero Gold libraries exhibited the lowest levels of rRNA, our method and the Adiconis method substantially depleted rRNA resulting in levels comparable to RiboZero Gold (Figure 4A). Of the libraries prepared with the commercially available kits, NEBNext and RiboErase had higher levels of rRNA. Gene expression estimates were highly correlated between replicate libraries within a given method and particularly high for the three methods with the most effective rRNA depletion (Figure 4B). The substantial amount of rRNA in the NEBNext and RiboErase samples combined with relatively shallow sequencing (3 million reads) was likely responsible for the lower correlation. Gene expression estimates calculated from our method correlated strongly with those calculated from both the Adiconis and RiboZero Gold kit (Figure 4C). While on-target depletion levels and the majority of gene expression levels are similar between our method and RiboZero rRNA depletion, a number of genes (108) are significantly differentially depleted in the RiboZero method (Figure 4D).

**Fig. 4.**
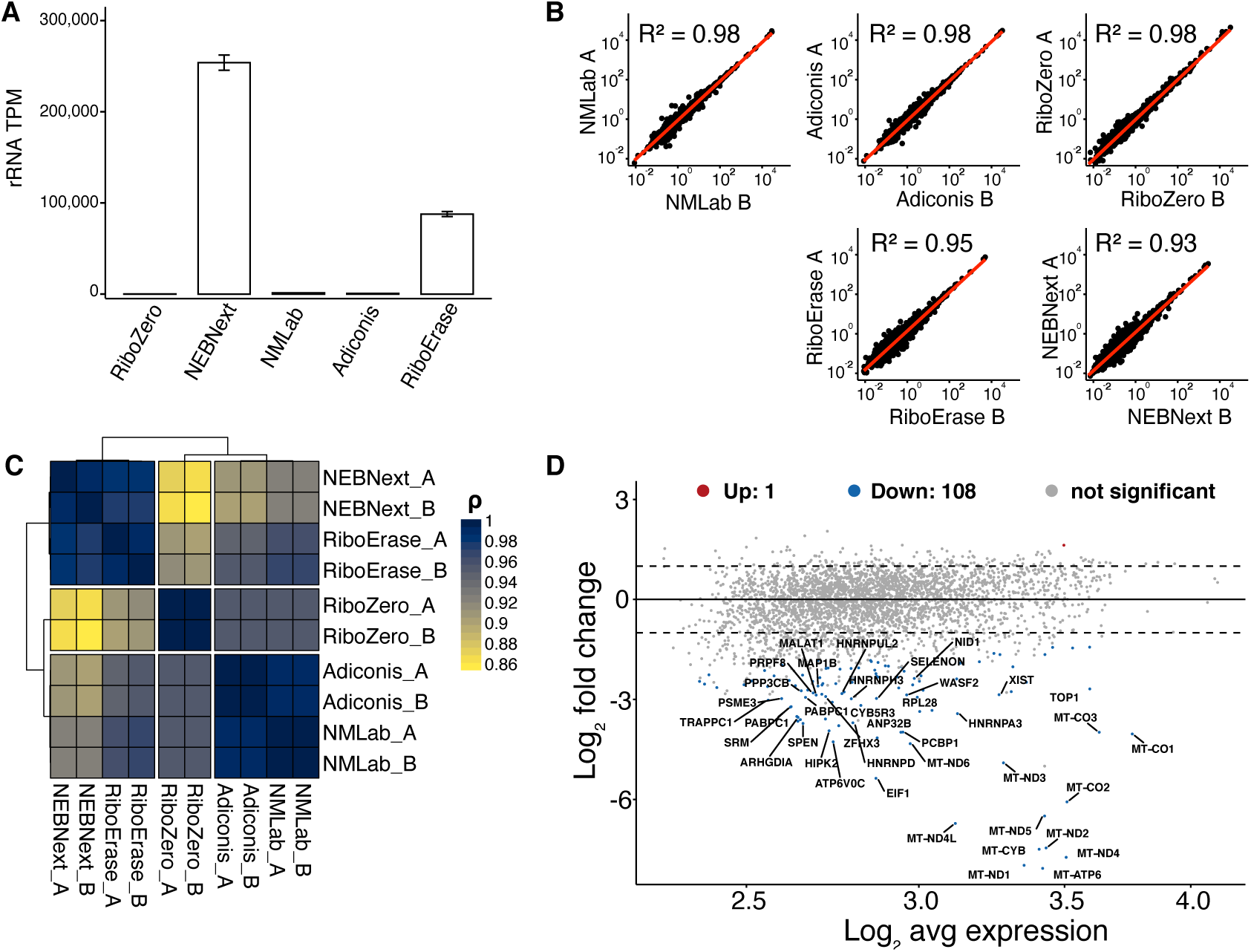
Comparison of five different rRNA depletion methods by RNA-seq. A) Transcripts per million (y-axis) of rRNA (including 5S, 5.8S, 18S, 28S, ITS, and ETS) for a given rRNA depletion method with error bars (standard error) for duplicates. B) Scatterplot of log10-transformed gene expression estimates with R-squared values comparing duplicate libraries for each rRNA depletion method. C) Heatmap of samples clustered by pearson correlation coefficients of gene expression comparing replicates of each rRNA depletion method. D) MA plot comparing average gene expression (y-axis) to differential gene expression (x-axis) of rRNA depletion versus Illumina RiboZero Gold (H/M/R) method colorcoded by statistical significance. Genes color-coded by statistical significance (higher in RiboZero (red), lower in RiboZero (blue), and no significant change (grey).

Closer examination of these RiboZero-depleted transcripts revealed mitochondrially-encoded mRNAs, and mRNAs encoding RNA-binding proteins, neither of which are described as targets for depletion by RiboZero (Supplemental Table 1).

### Efficient rRNA depletion for low input RNA

Having established our method was comparable to the standard in the field, RiboZero Gold, we next examined the effect and interaction between using either equimolar (1:1) or excess DNA oligos (5:1), and low (50 ng) or high (1 µg) input RNA. For both high and low RNA input amounts, rRNA depletion was more effective at the 5:1 DNA oligo to input RNA ratio. (Figure 5A). More effective depletion with higher DNA oligo:RNA input was particularly prominent for the low input sample. All triplicates exhibited a very high correlation of gene expression levels (Figure 5B). The correlation was strongest (R >.98) for the 5:1 oligo:RNA ratio regardless of the amount of input RNA.. Altogether, our data demonstrate that a 5:1 excess of DNA oligos produced the most efficient and consistent rRNA depletion. Furthermore, the method exhibited nearly identical gene expression estimates across nearly two orders of magnitude of input RNA.

**Fig. 5.**
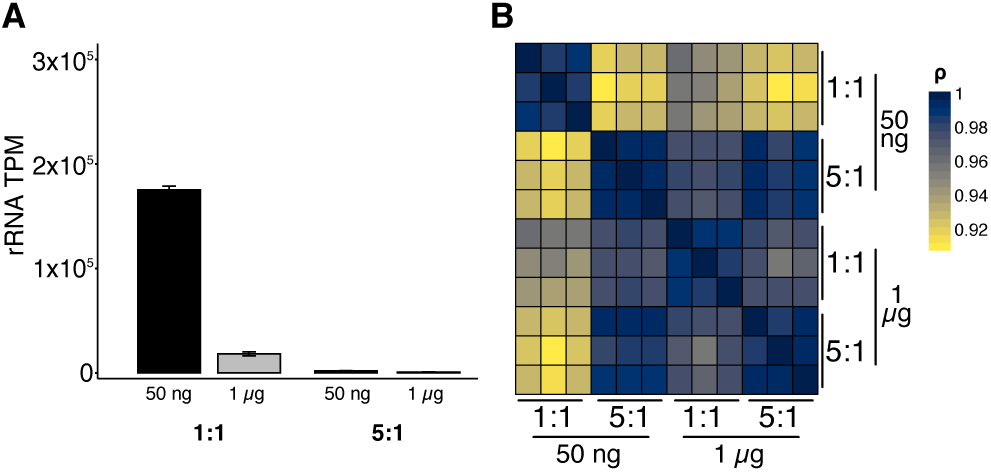
Comparison of oligo:RNA ratio for high and low input RNA by RNA-seq. A) Transcripts per million of rRNA (including 5S, 5.8S, 18S, and 28S) for a given rRNA input and oligo:RNA ratio with error bars (standard error) for triplicates. B) Heatmap of pearson correlation coefficients for gene expression across different rRNA input and oligo:RNA ratios for all replicates.

## Discussion

Through careful characterization of many parameters we have generated a streamlined, cost-effective, and robust method for rRNA depletion from human total RNA samples. The method can be performed with common lab equipment and supplies in just under two hours, with much of this time consisting of hands-off incubation steps (supplemental methods). We found that the standard RNaseH enzyme from E. coli underperforms in the context of rRNA depletion, likely due to the low reaction temperatures required for the enzyme. The 37°C temperature likely allows non-specific binding of the DNA oligos, forming RNA:DNA duplexes with non-target transcripts. This probable lack of specific binding in the context of E. coli RNaseH reaction temperature likely results in off-target depletion we observed. We found that thermostable RNaseH from the extremophile Thermus thermophilus is key for specific depletion of rRNA in RNA:DNA hybrids due to its intrinsic activity at higher temperatures. As the 50-mer DNA oligos used in this method have an average melting temperature of 73°C, the higher incubation temperature allowed by the use of a more thermostable RNaseH enzyme is crucial for allowing more specific binding of RNA to DNA and therefore less off-target depletion. This cost per reaction for rRNA depletion using our method is considerably less than available kits (Table 1). Since the RNAseH enzyme comprises the majority of the cost per reaction, moving forward it would be useful to explore alternative commercially produced thermostable RNaseH enzymes.

While our method is built on the foundation of previously published RNaseH-mediated depletion protocols (Adiconis et al., 2013; Morlan et al., 2012), we have made key improvements through careful optimization such as elevated reaction temperature (65°C compared to 45°C in Adiconis et al. 2013) discussed earlier. We also found that decreasing the incubation time to 10 minutes led to a decrease in off-target depletion while maintaining efficient rRNA depletion. We demonstrated that the protocol is suitable for a wide range of total RNA input. We successfully depleted rRNA from human cell line samples ranging from as low as 50ng to 1ug total RNA input. We have optimized the oligo:RNA concentration used in the reaction with oligos in molar excess (5:1, adding negligible additional cost for excess oligos) to guarantee efficient depletion of target rRNA. Other research groups have concluded this ratio to be optimal for RNaseH-mediated rRNA depletion in bacterial species (Huang et al., 2020). Subtractive hybridization and RNaseH-based cleavage of rRNA remain the predominant methods for rRNA depletion. A number of creative rRNA depletion methods have been performed but may be cumbersome and less efficient (Archer et al., 2014; Arnaud et al., 2016; Fauver et al., 2018; O’Neil et al., 2013; Zeng et al., 2020). Additional new commercial technologies have recently become available, such as QIAGEN’s FastSelect kit and Nugen AnyDeplete, which we have not tested in this study due to cost. Overall, the method we present has the optimal balance of cost and simplicity.

Our protocol has flexibility in both total RNA input and the oligos used for target depletion. While we used oligos designed for human samples from Adiconis et al. 2013, 50-mer reverse complementary oligos can easily be designed for other rRNA species, or pan-species targeting oligo pools, utilizing available published methods (Culviner et al., 2020). Additional spike-in oligos can be designed to target and deplete unwanted highly abundant transcripts such as globin mRNA for blood samples or the constitutively expressed transcript, 7SK. Unlike most commercially available kits, our streamlined protocol is not only flexible but transparent. The reaction compositions and oligo sequences are open source and should not result in depletion of unadvertised transcripts as was observed with RiboZero Gold. While this method can be used for many applications, it may not work well for rRNA depletion of ribosome profiling (Ribo-Seq) samples. Our pilot studies applying this method to Ribo-Seq samples initially looked promising but the oligo amount required for depletion was too much to be completely removed by DNase digestion (Supplemental Figure 2). Whereas small RNAs are depleted from general whole transcriptome RNAseq library preps during standard cleanups and size selections, short RNaseH degradation products could be problematic for RiboSeq as short products may be cloned into the library. Additionally, RNaseH-mediated approaches may be problematic for applications that need precise nucleotide resolution like Ribo-Seq (Zinshteyn et al., 2020).

This method is clearly valuable for standard whole transcriptome RNA-seq methods. This has implications for lowering the cost and/or improving mapping of mRNA modifications, capturing unstable precursor RNAs in metabolic labeling experiments, and any other application in which a few extremely abundant RNA species compromise sequencing depth. It can enable research surrounding less commonly used organisms without a well defined reference transcriptome, where some ribosomal RNA sequence (or one for closely related organisms) is known (Huang et al., 2020). RNaseH-mediated rRNA depletion has proven to be useful for samples with degraded input material such as archival clinical FFPE samples (Adiconis et al., 2013; Morlan et al., 2012), and for species that lack polyadenylated mRNA like bacterial transcriptomes. Our rRNA depletion method is applicable to many transcriptomic studies and is flexible, costeffective, and efficient.

## Methods

### Oligos

#### rRNA depletion oligos

50-mer oligos reverse complementary to Homo sapiens cytoplasmic and mitochondrial rRNA were ordered resuspended at 100µM from Integrated DNA Technologies as described in Adiconis et al. 2013. The oligos were pooled equimolarly and aliquoted for storage at -20°C.

#### qPCR oligos

Oligos for qPCR (supplemental methods) were designed to genes listed in the supplemental methods in the human transcriptome spanning exon-exon junctions, with an average melting temperature of 60°C and amplicon length of 100bp.

### Cell culture conditions

H295R cells were cultured on 10 cm plates in complete media (DMEM/F12, 10% cosmic calf serum, 1% insulin-transferrin-selenium) under 5% CO_2_ at 37°C. HEK293 Flp-In TREx FLAG-EGFP cells were cultured on 10 cm plates in complete media (DMEM, 10% fetal bovine serum) at 5% CO_2_ at 37°C.

### RNA extraction

All RNA used in experiments was collected from human cell lines above. RNA was harvested according to manufacturer’s instructions using the ZYMO Research Quick-RNA MiniPrep Plus kit and samples were treated with the provided on-column DNaseI treatment. RNA was quantified by Nanodrop OneC and Qubit RNA BR assay, and quality was checked by TapeStation RNA Screentape assay. Agarose gel electrophoresis was used to visually evaluate the quantity and integrity of rRNA. 2X RNA loading dye was added to samples at a final concentration of 1X, heated to 95°C for 5 minutes, then placed on ice for 2 minutes. The denatured RNA samples were loaded onto a non-denaturing 1.2% 1X TAE agarose gel and run at 100V for 3 hours. The gel was stained in 1X SYBRGold for 30 minutes then imaged with an EpiBlue channel on the Azure C150 imager.

### RNAseH-based rRNA depletion

#### Oligo:RNA hybridization

Total RNA was hybridized to rRNA depletion DNA oligos in oligo:RNA ratios described in the figures above and supplemental methods. The hybridization was performed in the 1X buffer of the RNaseH enzyme indicated (without MgCl_2_) in the presence of 50µM EDTA. The reactions were incubated at 95°C for 3 minutes to denature RNA, followed by a slow-ramp of -0.1°C/second to the temperature for RNaseH treatment as indicated for each experimental figure. Reactions were held at that temperature for 5 minutes then spun down briefly before immediately continuing to RNaseH treatment. Minus oligo control reactions contained nuclease free water instead of oligos.

#### RNaseH treatment

RNaseH enzyme in 1X buffer with necessary final concentration of MgCl_2_ was added to the hybridization reactions at the temperature and incubated for the time indicated in the experimental figures. “No RNaseH” reactions contain nuclease-free water instead of enzyme.

#### Cleanup

After RNaseH treatment, some samples were purified with an Agencourt RNAClean XP bead cleanup as detailed in the manufacturer’s instructions. With or without post-RNaseH cleanup, samples were treated with DNase enzyme in 1X DNase buffer to deplete leftover oligos from the reaction as outlined in the supplemental methods. DNase reactions were further cleaned with either the ZYMO RNA Clean and Concentrator-5 kit or RNAClean XP beads as detailed by the manufacturer’s instructions and supplemental methods.

### RT-qPCR

We were careful to use exactly the same volumes of RNA and cDNA for each qRT-PCR reaction. cDNA, minus RT, and no-template RT control reactions were generated using the Bio-Rad iScript cDNA synthesis kit following the manufacturer’s instructions. Resulting reactions were diluted with nuclease free water and used as input into qPCR reactions with a final concentration of 1X Bio-Rad iTaq Universal SYBR Green Supermix and 500nM primers listed in the supplemental methods. qPCR reactions were performed in triplicate for each sample and primer pair combination, including minus RT and no-template RT control reactions. Reactions were run on a Bio-Rad CFX 384 qPCR instrument. We used a deltaCt approach to compare levels of rRNA and housekeeping mRNA across samples. Ct values for all genes were normalized to their respective Ct values in the input RNA sample.

### Cost calculation

Cost per rRNA depletion method was calculated using the list price of the kit or necessary reagents when experiments were performed. Prices were calculated on a per reaction amount.

### RNA-seq

#### rRNA depletion

For each of the rRNA depletion methods tested, technical replicates were performed as described in the experiment figures and supplemental methods (including minus oligos and minus RNaseH controls if applicable) with total RNA from the same starting pool as input. In our home-brew method, 1µg total RNA was hybridized with 0.5µg rRNA depletion oligos in a 1:2 oligo:RNA concentration in 1X Lucigen Thermostable Hybridase RNaseH buffer (minus MgCl_2_) and 50uM EDTA by incubating at 95°C for 3 minutes followed by a slow ramp at rate of -0.1°C/second to 65°C. The hybridization reaction was held at 65°C for 5 minutes then treated with Lucigen Hybridase Thermostable RNaseH in 1X Lucigen Hybridase Thermostable buffer with a final concentration of 10mM MgCl_2_ at 65°C for 10 minutes. RNaseH reactions were purified with RNAClean XP beads according to the manufacturer’s instructions and then treated with Ambion Turbo DNase for 30 minutes at 37°C in 1X Turbo DNase buffer. The reactions were cleaned again with RNAClean XP beads and resulting RNA was eluted into nuclease free water. This method was also performed using 1µg and 50ng total RNA as input and hybridized at a 1:1 or 5:1 oligo:RNA ratio for rRNA depletion in triplicate, including controls. In the Adiconis 2013 method, total RNA was hybridized with 1µg of rRNA depletion oligos (1:1 oligo:RNA concentration) as described in Adiconis, 2013. The hybridization reaction, cleanup, DNase treatment, and final bead cleanup were also performed as described. The resulting bead cleaned RNA was eluted in nuclease free water. Total RNA was depleted of rRNA following manufacturer’s instructions for the following kits: Illumina, RiboZero Gold H/M/R; New England Biolab, NEBNext rRNA Depletion (human/mouse/rat) kit; KAPA Biosystems, RiboErase kit. The resulting depletion reactions were treated with DNase (if applicable) and cleaned with RNAClean XP beads as detailed in the manufacturer’s instructions for the respective depletion kit.

#### Library preparation

Each rRNA depletion sample for each method and replicate was used as input into the KAPA RNA HyperPrep library preparation kit. The sample was mixed with 2X Fragment, Prime, and Elute buffer, then incubated at 94°C for 4 minutes to fragment the samples. Library preparation was carried out according to the manufacturer’s instructions, using 5µL of a unique 7µM KAPA Dual-indexed adapter for each 1000ng starting input sample and 5µL of 1.5µM adapter for each 50ng starting input sample at the ligation step. PCR was performed with cycle number indicated in the supplemental methods for each sample followed by a cleanup and selection with the provided KAPA beads. The final libraries were eluted into 10mM Tris HCl, pH 8.0 and quantified using the Invitrogen Qubit dsDNA HS assay with Qubit 3.0 fluorometer. A portion of each library was diluted within the quantitative range and ran on the Tapestation 4200 with High Sensitivity D1000 Screentape assay.

#### RNA-Sequencing

Final libraries were submitted to the University of Colorado School of Medicine Genomics and Microarray Core for 150bp paired end sequencing on the Illumina NovaSeq 6000 platform. The libraries were sequenced to a median depth of 10 million paired end 150bp reads.

All transcript quantification was performed using Salmon version 0.13.1 (Patro et al., 2017) and the transcript index were built using Gencode v26 (https://www.gencodegenes.org/human/release_26.html).

All rRNA levels are expressed in transcripts per million (TPMs). After filtering for minimal detection thresholds, the pearson correlation coefficients of RNA expression between samples were calculated using R. Heatmaps of correlation coefficients were visualized using pheatmap version 1.0.12 (https://CRAN.R-project.org/package=pheatmap). Differential expression analysis was performed on rounded salmon counts using limma (Ritchie et al., 2015). Gene ontology analysis was performed using EnrichR version 2.1 (https://CRAN.R-project.org/package=enrichR). All R code used for analysis of qPCR and RNA-seq data, as well as, all qPCR and RNA-seq quantification data is available at https://github.com/mukherjeelab/rRNAdepletion. RNA-seq will be deposited in GEO and available at GSEXXXX.

## ACKNOWLEDGEMENTS

We thank members of the Mukherjee lab, David Bentley, Olivia Rissland, and Matt Taliaferro for discussions and advice and the Henriques lab for the bioRxiv template. N.M. acknowledges start-up funds from the RNA Bioscience Initiative, Boettcher Webb-Waring Biomedical Research Award (AWD-193075), and the Cancer League of Colorado (193481).

## AUTHOR CONTRIBUTIONS

A.B., A.R.M, and N.M. conceived the project; A.B. and N.M. developed the methodology; A.B. performed all experiments and collected all data; N.M. performed formal analysis and conducted the visualization; A.B. and N.M. wrote the original draft; A.B., A.R.M, and N.M. reviewed and edited the paper; N.M. acquired funding; N.M. provided resources; N.M. supervised the project.

---

**Fig. S1.**
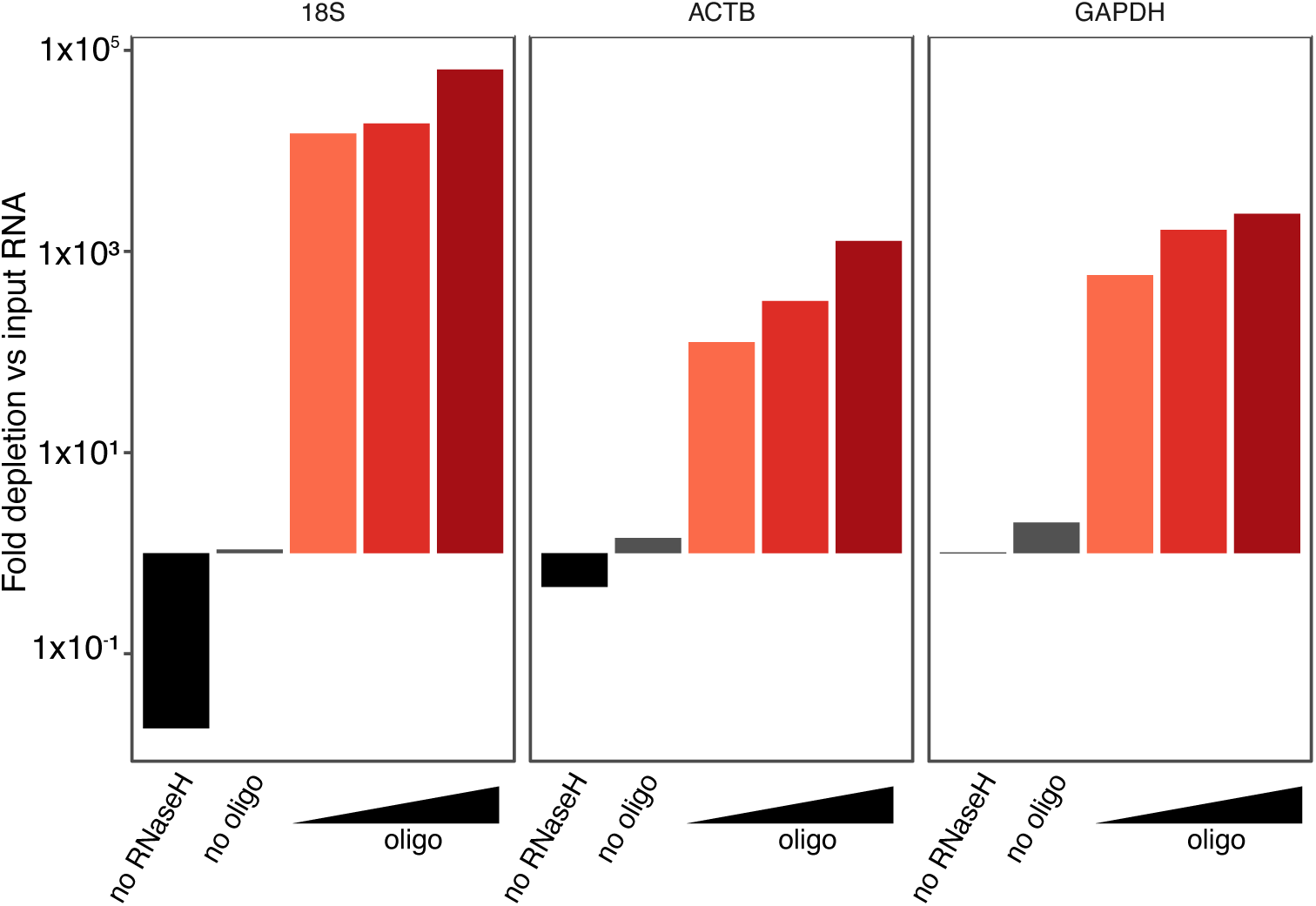
Control without RNaseH enzyme shows increase by qPCR likely due to priming of leftover DNA oligos. Fold depletion of 18S rRNA, ACTB, and GAPDH transcripts normalized to input total RNA by RT-qPCR performed on the RNA samples in Figure 2A. Total RNA samples incubated with oligos and DNase but without RNaseH (black bar), RNaseH and DNaseI without oligos (grey bar), or increasing oligo:RNA ratio (red bars) with RNaseH and DNaseI treatment. NEB E.coli RNaseH treatment for 1 hour at 37°C and NEB E.coli DNaseI treatment for 30 minutes at 37°C used for samples in this experiment.

**Fig. S2.**
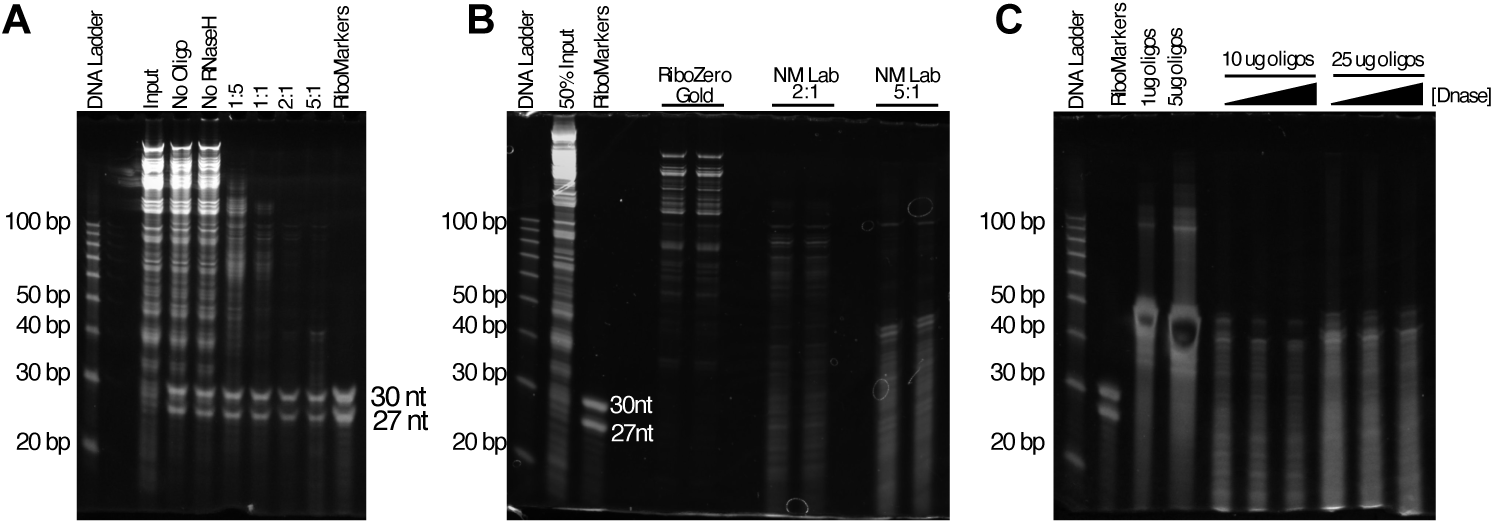
RNaseH-mediated rRNA depletion for Ribo-Seq. All samples were run on a 15% Denaturing PAGE gel. 30nt and 27nt RNA markes were spiked into input material. A) 1µg ribosome footprinting input material at various ratios of oligo:RNA concentration. B) 5µg ribosome footprinting input material depleted of rRNA using Illumina RiboZero Gold or NM Lab’s method at 2:1 and 5:1 oligo:RNA concentration. C) 10µg or 25µg rRNA depletion oligos treated with increasing con2entration of TurboDNase (5, 15, and 30U).

**Supplemental Table 1.** Gene ontology analysis of genes deferentially depleted by RiboZero Gold compared to our methodology.

| Term | Adj P-value | Genes |
| --- | --- | --- |
| mitochondrial inner membrane (GO:0005743) | 1.7e-07 | MT-ND6;MT-ND4;MT-ND5;MT-CO1;SDHC;MT-ND2;TYMS;MRPL12;MT-ND3;MT-ND1;MT-ATP6;MT-CO2;MT-CO3;MT-CYB |
| mitochondrial respiratory chain complex I (GO:0005747) | 2.4e-05 | MT-ND4L;MT-ND4;MT-ND5;MT-ND2;MT-ND3;MT-ND1 |
| cytoplasmic ribonucleoprotein granule (GO:0036464) | 1.7e-03 | LARP1;PCBP1;CNOT3;IGF2BP1;PABPC1;TOP1;RPL28 |

